# Hijacking and Integration of Algal Plastids and Mitochondria in a Polar Planktonic Host

**DOI:** 10.1101/2024.10.20.619283

**Authors:** Ananya Kedige Rao, Daniel Yee, Fabien Chevalier, Charlotte LeKieffre, Marie Pavie, Marine Olivetta, Omaya Dudin, Benoit Gallet, Elisabeth Hehenberger, Mehdi Seifi, Florian Jug, Joran Deschamps, Ting-Di Wu, Rebecca Gast, Pierre-Henri Jouneau, Johan Decelle

## Abstract

In oceanic plankton, various host organisms are capable of engulfing and temporarily integrating microalgae (photosymbiosis) or just their photosynthetic plastids (kleptoplastidy) as a solar-powered energy source. These cellular interactions can be considered to be representative of evolutionary steps in plastid acquisition in eukaryotes, but the underlying cellular mechanisms and dynamics are not fully understood. Here, we studied a kleptoplastidic dinoflagellate (RSD: Ross Sea Dinoflagellate), which is known to steal plastids of the microalga *Phaeocystis antarctica*. We tracked the morphology and activity of stolen plastids over several months by combining multimodal subcellular imaging and photophysiology. Upon integration inside a host vacuole, the volume of plastids and pyrenoids significantly increased and photosynthetic activity was boosted along with carbon fixation and transfer to the host. This may be supported by the retention of a 50-fold larger algal nucleus for ∼1 week. Once the algal nucleus was lost, there was a decrease in plastid volume and photosynthesis, but plastids were still beneficial for the host after > 2 months. Unlike other kleptoplastidic interactions, we showed that the algal mitochondrion was also stolen and retained for several months, transforming into an extensive network in close proximity with plastids. This highlights a new strategy in plankton along the continuum of plastid symbioses where both the energy-producing plastid and mitochondrion of a microalga are hijacked for several months by a host. This symbiosis that we found to be widely-distributed in polar regions suggests that plastid-mitochondrion interaction may have played a role in the evolution of plastid acquisition.

## INTRODUCTION

Marine plankton include a wide diversity of single-celled eukaryotes that can form complex cell-cell interactions during their life cycle [1,2]. Among the most intriguing examples are interactions between heterotrophic hosts that can engulf and temporarily integrate microalgae (photosymbiosis) or just their photosynthetic plastids (kleptoplastidy) as solar-powered carbon factories [2]. These plastid symbioses are widespread in the ocean and contribute significantly to biogeochemical cycles [56]. Photosymbiosis and kleptoplastidy are widely considered to represent fundamental evolutionary steps in the origin and distribution of plastids in eukaryotes [3,4]. Despite the ecological and evolutionary importance of photosymbiosis and kleptoplastidy, the structural and metabolic integration of a photosynthetic machinery into a host remains poorly understood. This may be explained by the fact that plastid integration in hosts is highly dynamic and can change over time. Therefore, in order to identify the nature and functioning of complex cellular interactions, it is crucial to acquire quantitative data at relevant spatial and temporal resolutions by integrating advanced high-resolution imaging.

Kleptoplastidy has been observed in multicellular organisms (sea slugs [5], acoels [6]), and protists (ciliates [7], dinoflagellates [8,9], Euglenozoa [10], Foraminifera [11], centrohelids [12]) in coastal and oceanic waters. A wide diversity of micro- and macro-algae species can be the source of plastids, but hosts typically rely on specific algae as prey. For instance, dinoflagellate hosts can acquire plastids from specific cryptophytes [14], diatoms [19] or haptophytes [9]. The most intriguing aspect of kleptoplastidy is how stolen plastids function in a host without their original nucleus for periods lasting from days to weeks [13]. In several kleptoplastidic interactions, stolen plastids have been reported to benefit to the host, providing photosynthetically-derived carbon energy (sugars and lipids), as well as nitrogen and sulfur compounds, and vitamins [10, 16]. In some cases, dinoflagellate and ciliate hosts can also steal and sequester the nucleus (called a kleptokaryon) and mitochondrion of the algal prey for several days [17–19]. When the algal nucleus is retained, stolen plastids can increase in size, divide and photoacclimate, but degrade after the nucleus is lost [18, 42 20,60–61]. The mechanisms by which stolen plastids morphologically and physiologically adapt within a foreign host over time and the underlying molecular processes regulating functional stolen plastids remain unresolved. Exploring the cellular mechanisms of kleptoplastidy in protists can provide critical insights into the evolutionary acquisition of plastids in eukaryotes.

A kleptoplastidic dinoflagellate species found in Antarctic waters (Ross Sea Dinoflagellate, RSD) steals plastids from the bloom-forming haptophyte microalga *Phaeocystis antarctica* [9, 21, 22]. This dinoflagellate is part of the Kareniaceae family, most members of which harbor permanent plastids derived (*via* tertiary endosymbiosis acquisition) from haptophytes [15]. The RSD host strictly depends on kleptoplastidy for its growth, most likely by benefiting from photosynthetic products. In laboratory culture, stolen plastids have been reported to remain for over two years inside hosts [21]. This capacity to retain plastids for extended periods could be explained by the genetic toolkit of the host, composed of nucleus-encoded photosynthesis genes acquired from haptophyte and other microalgal sources (Horizontal Gene Transfer, HGT) [22]. This includes certain genes from Photosystem I and the cytochrome b6/f complex with specific sequences to target proteins to the stolen plastid. However, some essential nucleus-encoded genes, such as photosystem II (PsbO), are missing, which should dramatically limit photosynthetic activity of stolen plastids over time. A potential dependence of photosynthesis on PSI through Cyclic Electron Transport has therefore been proposed [22, 25]. Fundamental information on this kleptoplastidy is lacking, particularly concerning the dynamics of the interaction and the fate of stolen plastids over time and whether other algal organelles are retained in the host.

In this study, we used the Ross Sea dinoflagellate to investigate how stolen plastids are integrated and maintained within a host at relevant temporal and spatial scales. We employed an innovative combination of multimodal imaging and molecular tools, including volume electron microscopy for three-dimensional (3D) reconstruction of plastid morphology, Nano-scale Secondary Ion Mass Spectrometry (NanoSIMS) to track carbon fluxes, and photophysiology to monitor the photosynthetic activity of stolen plastids over several months. This multiscale study revealed a new strategy in protists whereby the host hijacks not only the plastids, but also the mitochondria of microalgae, both of which remain functional for several months without their original nucleus. These closely interacting stolen organelles were morphologically and metabolically remodeled in the host, with plastids notably exhibiting increased volume and boosted photosynthetic activity during the first two weeks. This study improves our knowledge on plastid symbioses in marine ecosystems and provides insights into the potentially fundamental role of the mitochondrion during the evolutionary acquisition of plastids in eukaryotes.

## RESULTS AND DISCUSSION

### Enlargement and enhanced photosynthetic activity of newly integrated stolen plastids

The subcellular organization of the kleptoplastidic association between the microalga *Phaeocystis antarctica* and the dinoflagellate host (RSD), with a focus on the morphology of stolen plastids, was first investigated by multiscale imaging in the presence of algal prey (fed conditions, Figure 1A). Confocal fluorescence microscopy revealed that host cells contained numerous stolen plastids (n=∼40 per cell), which appeared to be larger in size compared to native plastids of the prey (Figures 1B and 1C). 3D reconstruction and volumetric analyses based on FIB-SEM (Focused-Ion Beam Scanning Electron Microscopy) images, conducted from three host cells and eight prey cells, revealed that the volume of stolen plastids was 6 times greater than that of native plastids of the algal prey (11.11 µm^3^ ± 3.30, n=14 vs 1.91 µm^3^ ± 0.43, n=12, respectively), and 3 times greater than that of plastids from dividing algal prey (3.50 µm^3^ ± 0.62, n=9) (Figures 1F and 2A-C; Table S1). Stolen plastids were not only larger but also contained 12 times more chlorophyll (1.24 ng ± 0.17 per stolen plastid vs 0.11 ng ± 0.014 per native plastid, Table S1) and contained more thylakoid membranes, which sustain the light reactions of photosynthesis (Figures 1D and 1E). More specifically, the ratio of thylakoids to the total surface area was higher in stolen plastids (0.71 ± 0.05, n = 9) than in the prey (0.57 ± 0.09, n = 18) (Table S1). Some of the stolen plastids that were observed were seemingly undergoing division based on their large volumes and possible constriction sites with a continuous pyrenoid (Figure 2B). FIB-SEM images also revealed that stolen plastids were enclosed within a single large vacuole (hereafter called symbiosome) (Figure 1E).

**Figure 1.**
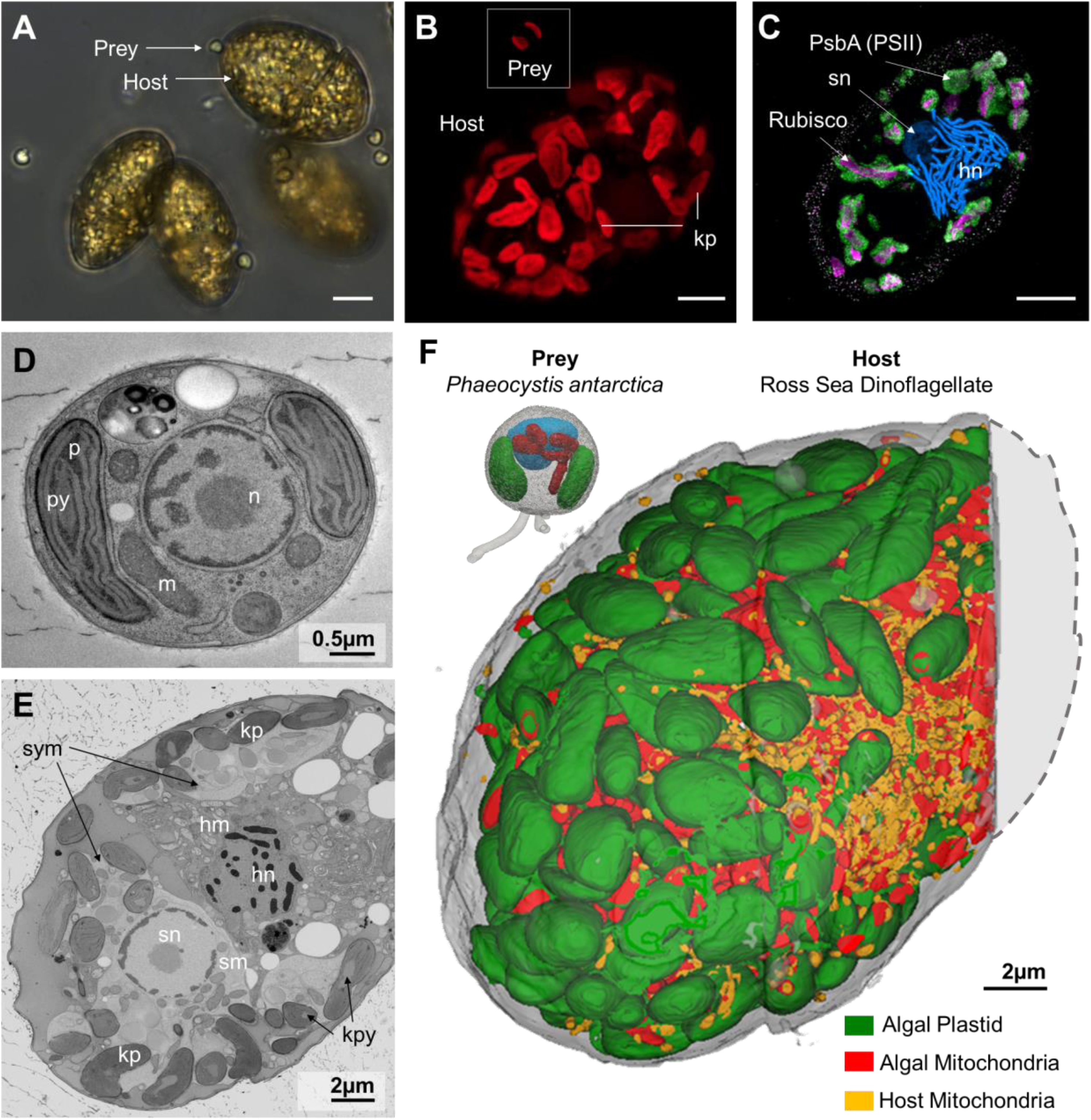
Subcellular organization of the kleptoplastidy between the host dinoflagellate (Ross Sea Dinoflagellate; fed condition) and its microalga prey *Phaeocystis antarctica* unveiled by multimodal imaging. **(A)** Light microscopy image of the co-culture of the Ross Sea Dinoflagellate (RSD, Kareniaceae; 20-40 µm) and its microalgal prey *Phaeocystis antarctica* (haptophyte; 2-5 µm). **(B)** Confocal fluorescence images showing the morphology of plastids unveiled by the red chlorophyll fluorescence in the microalgal *Phaeocystis* (prey, in the box) and in the host dinoflagellate (kleptoplast, kp). Up to 40 stolen plastids could be observed in the host and tended to be larger than native plastids in the algal prey. **(C)** Confocal fluorescence image of an expanded host cell after Ultrastructural Expansion Microscopy (UExM), stained using PSII PsbA antibody (thylakoids; green), RuBisCO antibody (pyrenoid; magenta), and NIR694 Nuclear Dye (host nucleus (hn), stolen nucleus (sn); blue). This shows the presence of the algal nucleus at this stage (fed conditions). **(A-C)** Scale = 5 µm. **(D)** Transmission Electron Microscopy (TEM) image of the algal prey *P. antarctica* with its plastids (p), pyrenoid (py), mitochondria (m) and nucleus (n); Scale = 0.5 µm. **(E)** Micrograph from a Focused-Ion Beam Scanning EM (FIB-SEM) stack of a dinoflagellate host with its nucleus (hn) and mitochondria (hm), as well as the stolen organelles of the microalga *P. antarctica*: plastids (kp) and their pyrenoids (kpy), mitochondria (sm) and nucleus (sn). These stolen algal organelles were enclosed in a large compartment, (symbiosome; sym); Scale = 2 µm. **(F)** Three-dimensional reconstruction based on FIB-SEM of the microalga *P. antarctica* (top left; prey) with its plastids (green), mitochondria (red) and nucleus (blue), and the subcellular organization of the RSD host in fed condition with stolen plastids (green) and mitochondria (red), and host mitochondria (orange) (based on a stack of ∼2800 images, 6 nm resolution). 3D reconstruction confirmed the significant enlargement of stolen plastids compared to their native state. The outline of the host cell was completed with a dashed line; Scale = 2 µm.

**Figure 2:**
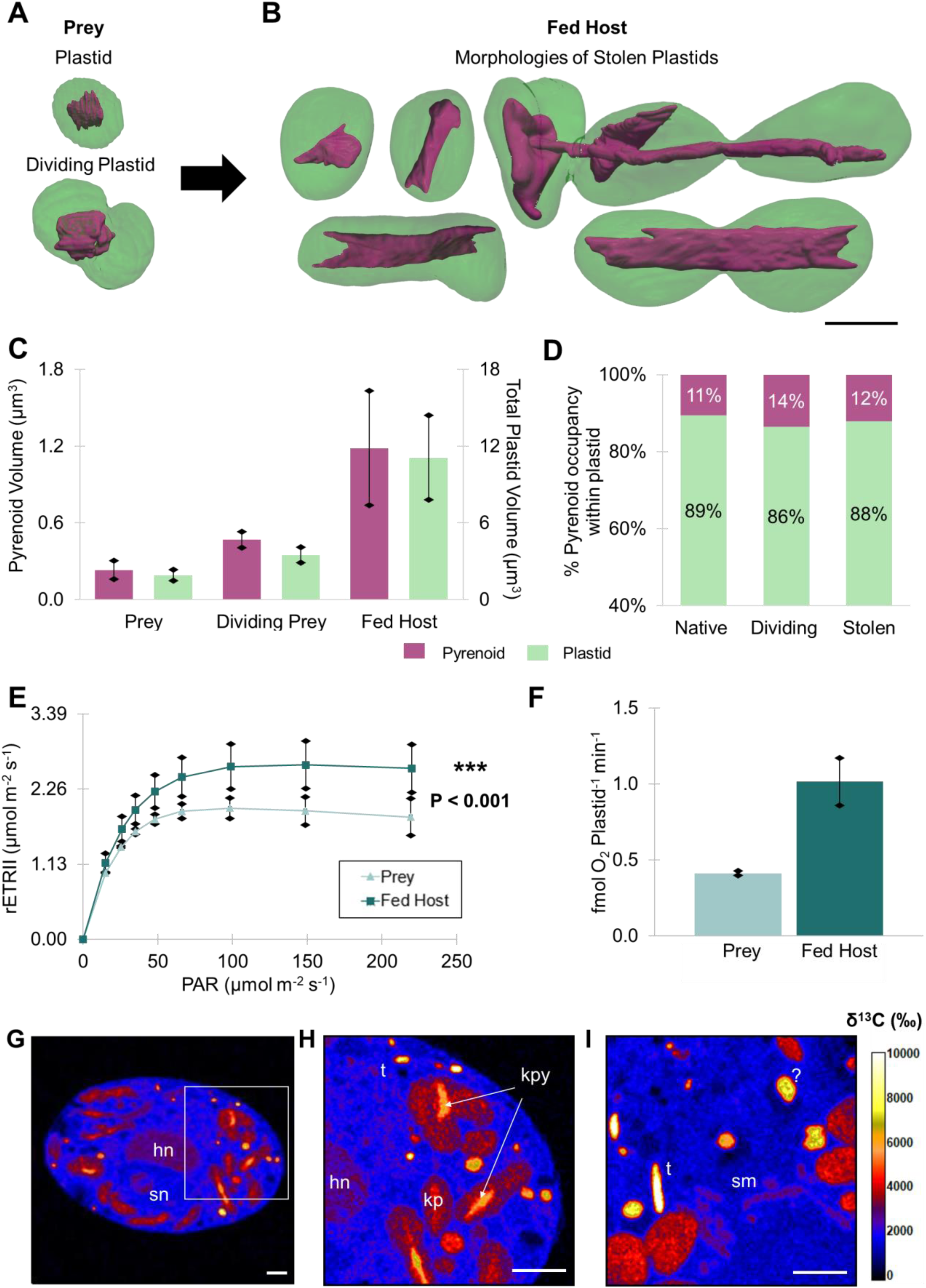
Morphometrics and photophysiology of stolen plastids within the dinoflagellate host (fed condition). (A-B) FIB-SEM based 3D reconstructions of native plastids (green) from non-dividing and dividing algal prey *P. antarctica* (A), and stolen plastids (green) in the dinoflagellate host (RSD) (B), including their carbon-fixing pyrenoid (magenta). Stolen plastids displayed different morphologies and some of them seemed to divide as shown by the constriction point and continuous pyrenoid (observations done on three different host cells); Scale = 2 µm. **C)** Volume of individual plastids (green; right axis) and their pyrenoids (magenta; left axis) in native (non-dividing microalga, n = 12 and dividing microalga, n = 9) and stolen states (fed host; n= 14) assessed from 3D models (µm^3^). On average, volumes of stolen plastids and pyrenoids increased six and five times, respectively, compared to native plastids. Data available in Table S1. **(D)** Occupancy (%) of the pyrenoid (magenta) within the native and stolen plastids (green) assessed from FIB-SEM reconstructions. **(E)** Photosynthetic efficiency was measured by the relative Electron Transport Rate through Photosystem II (rETR II, µmol m^-2^ s^-1^) from a culture of *P. antarctica* (triangle) and a culture of fed dinoflagellate hosts (square) over a range of eight light intensities from 15 to 219 µmol photons m^-2^ s^-1^ (n = 3; each measure was obtained from three culture replicates). ETR curves showed that photosynthetic efficiency was higher in stolen plastids compared to native plastids (P value < 0.001). **(F)** Gross photosynthetic oxygen per plastid (fmol O_2_ plastid^-1^ min^-1^) in fed hosts was 2.5 times higher compared to the microalga (n = 3; each measure was obtained from three culture replicates). **(G-I)** NanoSIMS ^13^C/^12^C isotope ratio images of fed hosts from resin sections showing ^13^C enrichment in stolen algal organelles (plastids (kp) and their pyrenoids (kpy), mitochondria (sm), nucleus (sn)), as well as in the host nucleus (hn), trichocysts (t) and other unknown structures. **(H)** Zoom in of G. Overlays available in Figure S3. **(G-I)** Scale = 2 µm. See also Tables S1 and S2.

In order to evaluate how morphological changes of stolen plastids influence their physiology, photosynthetic activity was compared with native plastids of *P. antarctica*. The relative electron transfer rate through PSII (rETR II) was significantly higher (P < 0.001 (Table S2)) in stolen plastids in a range of light conditions from 15 to 219 µmol photons m^-2^ s^-1^ (Figure 2E). The alpha value from the ETR curve tended to be higher in stolen plastids, which could indicate a higher effective antenna size (Figure S5; Table S2). In addition, oxygen measurements showed that gross production per plastid was 2.5 times higher in stolen plastids compared to native ones in *P. antarctica* (1.01 fmol O_2_ plastid^-1^ min^-1^ ± 0.16, n = 3 and 0.41 fmol O_2_ plastid^-1^ min^-1^ ± 0.01, n = 3) (Figure 2F). Our results indicate that photosynthetic production of newly acquired plastids was enhanced in the new host, correlating with increased volume, chlorophyll content and thylakoid stacking. These fluorescence-based measurements on PSII suggest that the host RSD retains functioning plastids, conflicting with previous studies reporting that photosystem II (PSII) was absent or diminished in its function ([22] [25]). We further verified PSII activity with DCMU (3-(3,4-dichlorophenyl)-1,1-dimethylurea), an inhibitor of PSII, which caused a drop in Fv/FM (from 0.64 to 0.35) and no detectable ETR. Combining Western blot analyses and Utrastructure Expansion Microscopy (U-ExM; [63]), recently applied on various planktonic eukaryotes [64] [65], we validated the presence and localization of PsbA (plastid-encoded photosystem II subunit) in the thylakoids of stolen plastids (Figure 1C; Figures S1 and S2).

### Carbon fixation by stolen plastids and carbon transfer to the host cell

In order to further understand the function and potential benefits of the newly integrated plastids in host cells (fed condition), their capability for carbon fixation was also assessed. We first examined the morphology of the pyrenoid, the CO_2_-fixing compartment within the plastids, containing the enzyme Rubisco [28]. Pyrenoids in stolen plastids tended to change from a round to an elongated rod-like shape (Figures 2A and 2B) and their volume was 5-fold higher on average than pyrenoids in prey plastids (1.19 µm^3^ ± 0.45, n = 14 and 0.23 µm^3^ ± 0.07, n = 12; respectively) (Figure 2C; Table S1). Pyrenoids occupied a similar volume (11-12%) within stolen and native plastids, demonstrating that pyrenoid to plastid volume ratio is maintained in the enlarged stolen plastids (Figure 2D). UexM and Western blots indicated that the plastid-encoded enzyme Rubisco (RbcL, large subunit) was present and localized in pyrenoids of stolen plastids as expected (Figure 1C; Figures S1 and S2).

To assess whether stolen plastids can fix inorganic carbon and provide photosynthates to their new host, ^13^C isotope ratio images were obtained using NanoSIMS for several host cells incubated for 24 hours with labeled ^13^C-bicarbonate. NanoSIMS images revealed significant ^13^C enrichment in stolen plastids, especially in pyrenoids (6000-8000‰) and the thylakoid area (4000‰) (Figures 2G-I and S3). Significant ^13^C enrichment was also detected in the host cell, especially in trichocysts and several unidentified structures (4000-8000‰), as well as in the host nucleus (2100-4000‰), demonstrating that stolen plastids export carbon to the host. Thus, once engulfed in their new host, stolen plastids undergo significant volume expansion with higher photosynthetic activity, which is clearly beneficial to the host as a source of carbon. This raises the question about the underlying mechanisms of plastid remodeling and photosynthetic enhancement in this association.

### Retention and enlargement of the algal mitochondrion and nucleus in the host

Electron microscopy revealed the presence of intact engulfed algal cells within fed hosts, indicating that plastid uptake takes place through phagotrophy of the entire algal cell (Figure S4). Upon engulfment, observations with expansion and electron microscopy revealed that the host not only retained several plastids but also the nucleus, mitochondria and possibly other organelles from the algae within a single vacuole (Figures 1 and 3, Figure S4). Of note, ^13^C enrichment was observed in these stolen organelles (2100-4000‰) by NanoSIMS, indicating that they utilize photosynthetically-derived carbon (Figures 2G-I; Figure S3). FIB-SEM imaging revealed that the algal mitochondrion dramatically expanded in volume (more than 20-fold increase, Table S1), and transformed into a highly reticulated network, in close association with plastids (Figures 1F and 3D). Mitochondrial protrusions into a “pocket-like” structure were observed within the plastid (Figures 3D and S4). A similar phenotypic transformation of the mitochondrion and close interaction with plastids has been reported in the microalga *Phaeocystis* in symbiosis within acantharian hosts [29]. This significant mitochondrial development in photosymbiosis and kleptoplastidy models indicates an integral role of the algal mitochondrion in photosynthesis and bioenergetics, as reported in diatoms and plants [31, 32].

**Figure 3:**
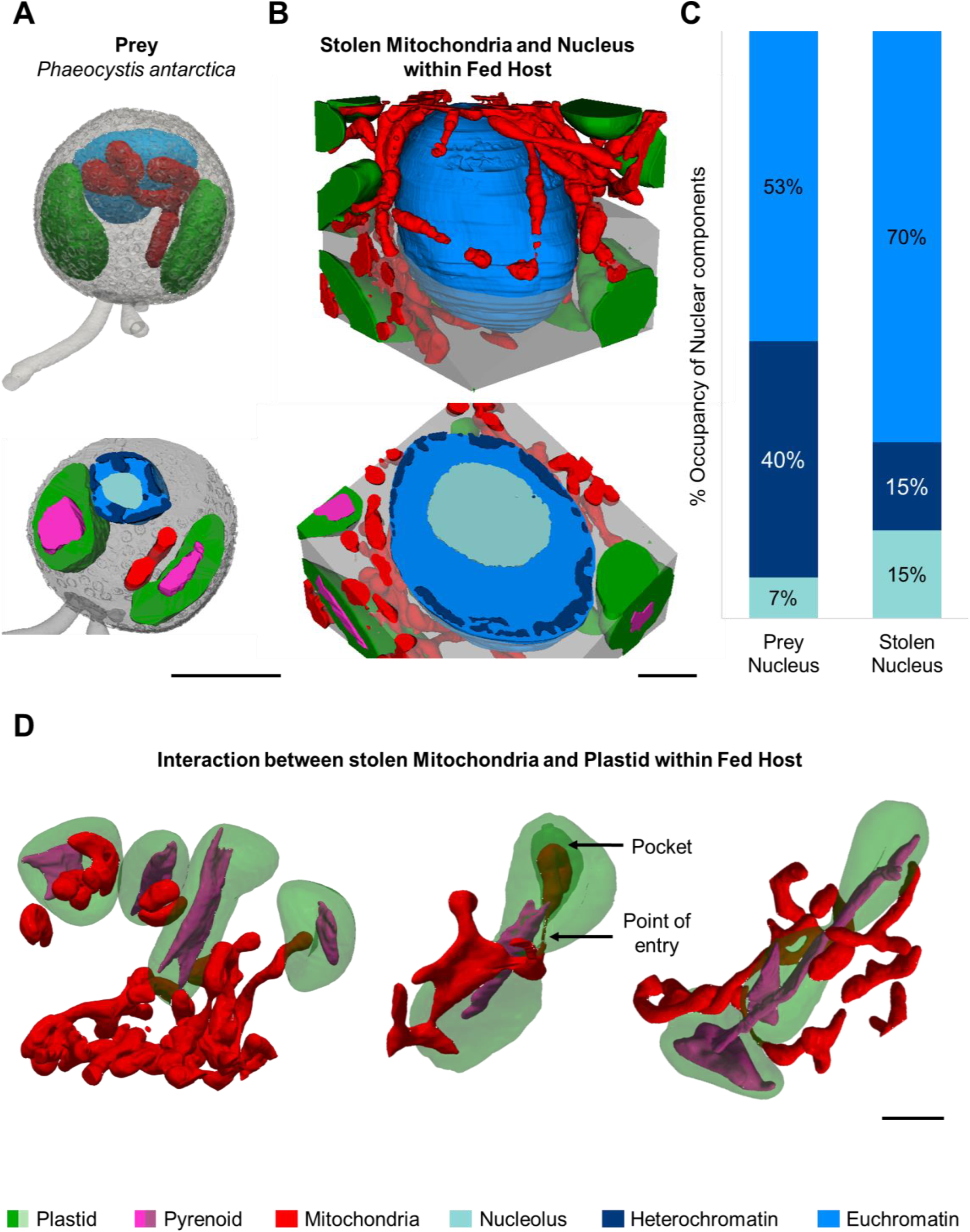
Three-dimensional reconstruction of newly stolen mitochondria and nucleus of the algal prey within the dinoflagellate host. **(A)** 3D reconstruction based on FIB-SEM of a *Phaeocystis antarctica* cell (prey) and a coronal section (below), with nucleus (blue), two plastids (green) and mitochondria (red). **(B)** 3D reconstruction of a host region focusing on the stolen nucleus and mitochondria of *P. antarctica*, and an axial section (below) showing the significant increase in volume of these algal organelles. Particularly, the algal mitochondria formed an extensive network around the plastids and algal nucleus. **(C)** Volume occupancy (%) of different nuclear compartments - euchromatin (blue), heterochromatin (dark blue, electron dense nuclear structures), and nucleolus (light blue) - within the volume of native and stolen prey nucleus (prey nucleus, n = 3; stolen nucleus, n = 1). This shows that heterochromatin volume occupancy decreased and the nucleolus volume occupancy increased in stolen nucleus. **(D)** 3D reconstructions of different interactions between the stolen mitochondria (red) and plastids (green) within the host. Black arrows indicate the site of entry of the mitochondrion into the plastid and a pocket-like structure showing close interaction between both organelles. **(A,B,D)** Scale = 2 µm. Plastid (green), pyrenoid (magenta), mitochondria (red), nucleolus (light blue), heterochromatin (dark blue), euchromatin (blue), rest of the cell/symbiosome (grey). See also Table S1.

In addition to the algal mitochondrion, one or several microalgal nuclei was/were observed in continuously-fed host cells (Figures 1C and 1E; Figures S1 and S4). Compared to the original nucleus of the microalga, the volume of stolen nucleus increased 50-fold, with a ∼90 and ∼15 fold increase in the volume of the nucleolus (ribosome factory; [33]) and heterochromatin, respectively (Figures 3A-C; Table S1). There was also a remodeling of the nuclear compartments, whereby heterochromatin occupancy (within total nucleus volume) decreased by 25% while nucleolus occupancy increased by 8% (Figure 3C), suggesting higher transcriptional activity in the algal nucleus. At this stage of the kleptoplastidy when prey is available, the presence of the algal nucleus and mitochondrion may support the growth and activity of stolen plastids by providing proteins and metabolites [18, 42].

### Tracking the morphology and activity of stolen plastids during several months of starvation

Whereas preceding observations were based on newly acquired plastids, in culture conditions where algal prey was continuously present, in the natural environment hosts may be temporarily starved when prey availability decreases (e.g. austral winter). In this kleptoplastidic association, a previous study reported that stolen plastids could be maintained for over two years in the absence of algal prey [9]. However, there is no information on the activity of stolen plastids (except for the absence of host nucleus-encoded PSII genes [22, 25]) and the fate of the algal mitochondrion and nucleus during starvation. In order to address this knowledge gap, a time-course experiment was conducted to monitor starved host cells in the absence of algal prey for up to 30 weeks (Figure 4). Physiological measurements and microscopy observations were carried out at different time points (weeks 1, 3, 9 and 30). After 1 week of starvation, electron microscopy observations showed that ∼2% of host cells still contained at least one algal nucleus (n= 53 cells), whereas none were present at week 3 and beyond. It is likely that algal nuclei rapidly disappear in most host cells by dilution over multiple host divisions. Because the nucleus of photosynthetic organisms is fundamental for the maintenance and physiology of plastids [34], its absence raises questions around how stolen plastids function and provide benefits for the host.

**Figure 4:**
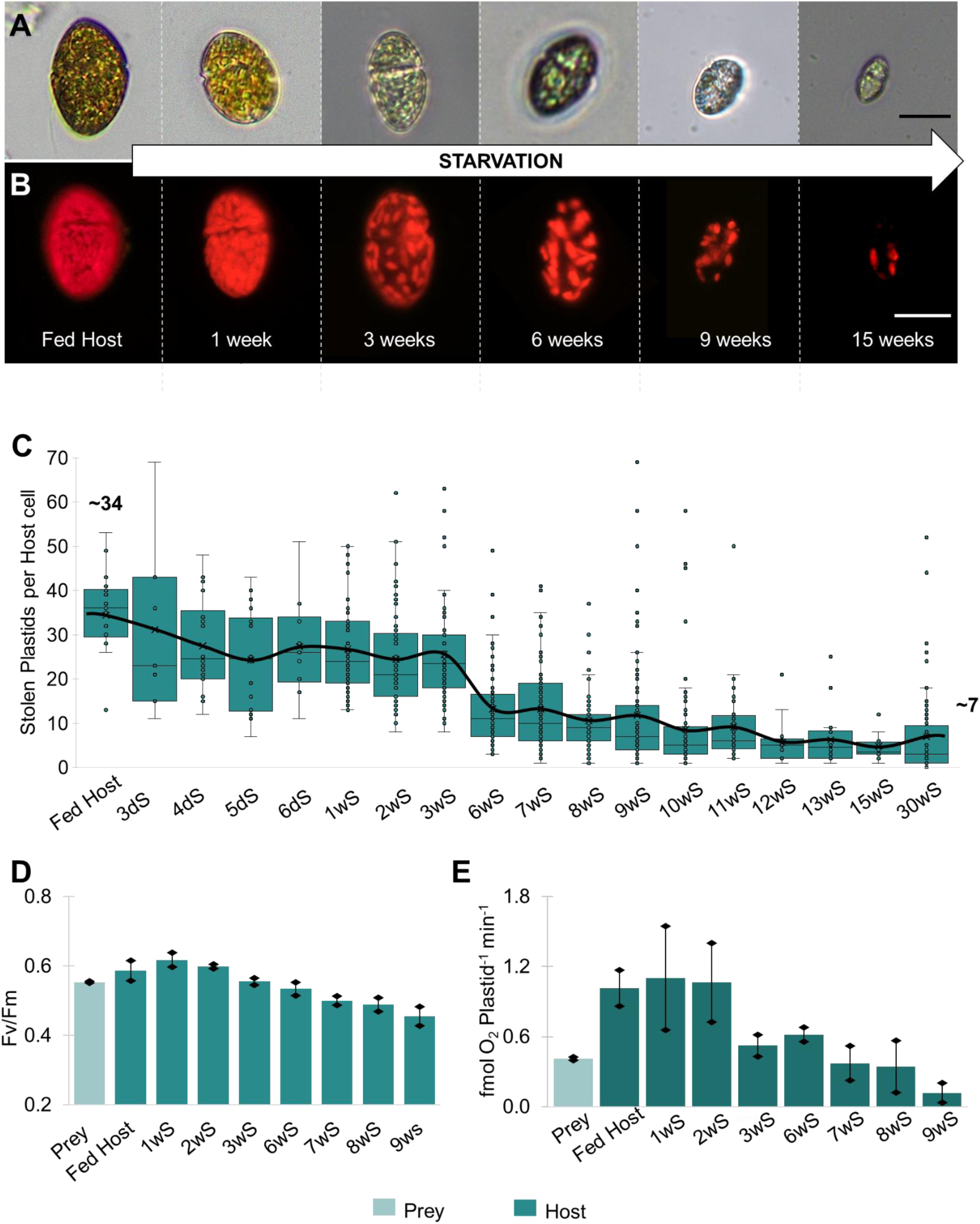
Retention time and photophysiology of stolen plastids in host cells during several weeks of starvation of (in the absence of algal prey) **(A-B)** Light microscopy images of host cells (RSD dinoflagellate) over 15 weeks of starvation in bright field (A) and fluorescence (B, red chlorophyll fluorescence); Scale = 20 µm. **(C)** Box plot showing the number of stolen plastids per host cell over 30 weeks of starvation (abbreviated as “wS”). Plastid counts were performed from fluorescence images taken at different time points from 18 to 88 host cells. At the beginning of starvation, the number of stolen plastids per host cell was around 34 and decreased to ∼ 7 at week 30. Plastid numbers were also assessed every day during the first week (3dS, 4dS, 5dS and 6dS). The black line connects the average plastid number of each week. Increased number of outliers were observed in the later weeks (> 3 weeks) of starvation and corresponded to very large host cells in a resting-like stage. **(D)** Bar plot of Maximum Quantum Yield (Fv/Fm) of Photosynthesis from a culture of microalga *Phaeocystis antarctica* (prey) and culture of fed and starved host cells over 9 weeks. Fv/Fm tended to slightly increase at week 1 (1wS, Fv/Fm = 0.618) compared to the fed host (Fv/Fm = 0.587), followed by a continuous decrease (Fv/Fm = 0.455 at week 9; 9wS) (n = 3; each measure was obtained from three culture replicates). **(E)** Gross oxygen production per plastid (fmol O_2_ Plastid^-1^ min^-1^) measured from the microalga *P. antarctica* (prey) and starved host cells over 9-weeks. Compared to newly stolen plastids (fed conditions), oxygen production per plastid was similar at week 1(1.10 fmol O_2_ Plastid ^-1^min^-1^ compared to 1.01 fmol O_2_ Plastid ^-1^min^-1^), and then gradually decreased over the next few weeks (0.12 fmol O_2_ Plastid ^-1^min^-1^ at week-9; 9wS) (n = each measure was obtained from three culture replicates). See also Table S2.

During the first week of starvation, dinoflagellate host cells divided (0.6 division/day on average), but plastid number per host remained relatively stable (∼30 on average; Figures 4A-C). This indicates that stolen plastids were able to divide during the first week, which was supported by our FIB-SEM observations (Figure 2B). Between week 1 and week 30, host cells decreased in size and gradually lost plastids (down to ∼7 on average; Figures 4A-C), indicating that plastids do not divide in the host once the algal nucleus is lost. On rare occasions, host cells with ∼40 plastids were still observed after 30 weeks of starvation, but the morphology of these cells suggests that they were resting stages.

Physiological activity and morphology of stolen plastids were then investigated during the starvation period. Fv/Fm decreased from 0.64 in “continuously-fed” condition to 0.46 at week 9 (Figure 4D). Photosynthetic oxygen production per plastid was relatively stable during the first two weeks of starvation (around 1 fmol O_2_ plastid^-1^ min^-1^, n = 3 culture replicates) and similar to that of freshly acquired plastids (Figure 4E). Subsequently, a continuous decrease in oxygen production per plastid was observed at week 3 (0.5 ± 0.09 fmol O_2_ plastid^-1^ min^-1^) and week 9 (0.12 ± 0.08 fmol O_2_ plastid^-1^ min^-1^). Corroborating these results, ETR measurements, including the alpha parameter, also indicated that PSII activity decreased during starvation (Figure S5). Yet, these measurements indicate that PSII was present and functional in stolen plastids for more than 9 weeks. Western blot analyses confirmed that subunits of PSII (PsbA) and PSI (PsaC) were still present during this starvation period (potentially still encoded by the stolen plastids or stable over time) (Figure S2). The nucleus-encoded PSII subunit PsbO was detected at week 3, but not at week 9 (likely below the detection limit due to the very low number of plastids/cell). We hypothesize that PSII starts to degrade at later stages of the kleptoplastidy, impairing the electron transport chain within stolen plastids as observed here.

The diminished photosynthetic activity of stolen plastids was also accompanied by drastic morphological changes (Figures 5A-B): their volume significantly decreased by 35% at week 9 (7.18 µm^3^ ± 2.72, n = 14) compared to freshly engulfed plastids, with much fewer thylakoid membranes and lower chlorophyll content (Figures 5 and S6; Table S1). Note that these plastids at week 9 were still larger than the native ones in *P. antarctica*. Old plastids also exhibited internal vacuoles/vesicles that seem to have be expelled into the symbiosome (Figures 5A and S6). This could be a sign of senescence in aging plastids (or gerontoplasts) where autophagic degradation can take place [35, 58–59].

**Figure 5.**
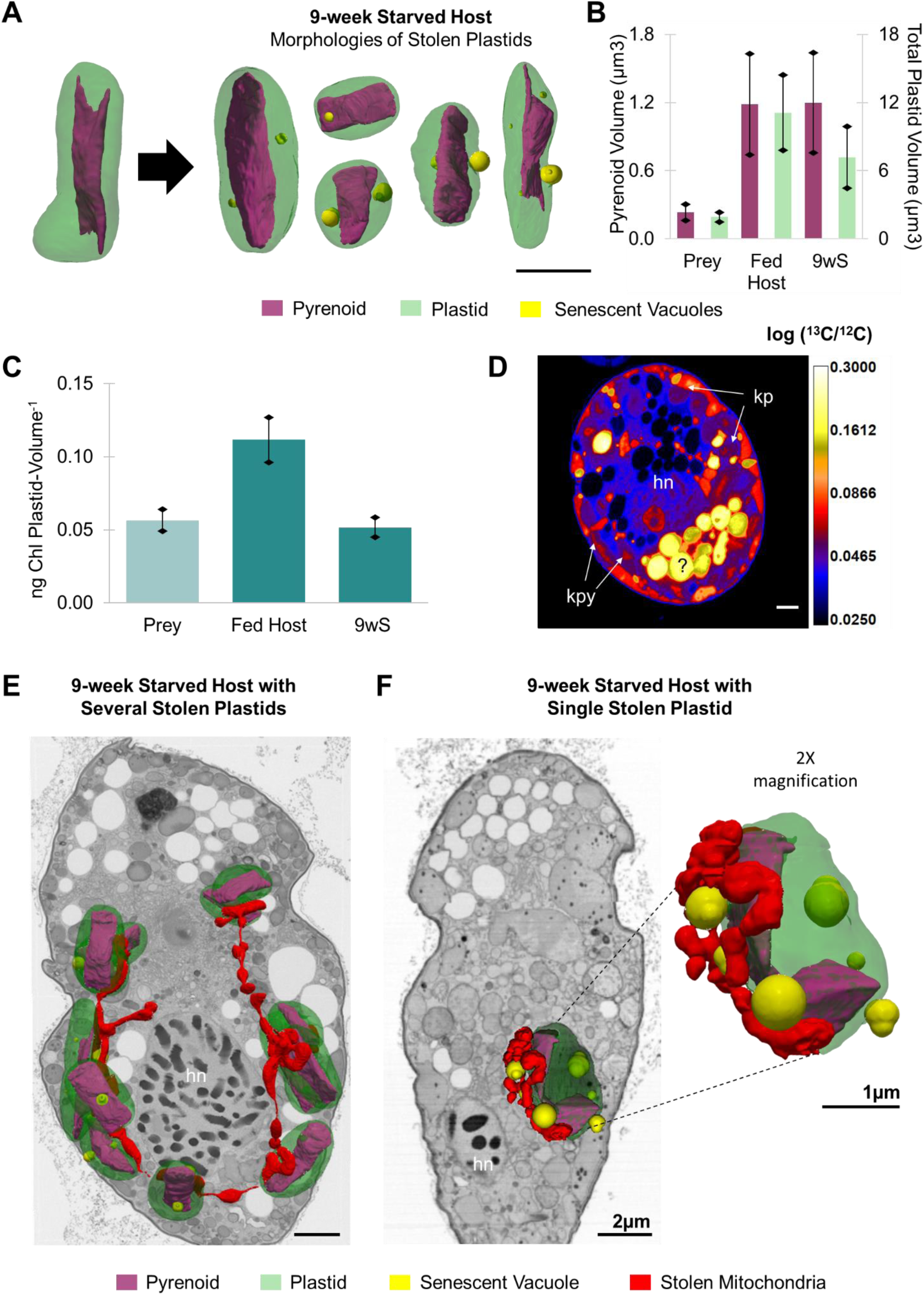
Morphology and physiology of “old” stolen organelles (the algal plastids and mitochondria) at week 9 of starvation. **(A)** 3D-reconstruction of stolen plastids (green) and their internal pyrenoid (magenta) from fed and starved (9 weeks) hosts. Several unknown spherical structures (yellow) were observed at week 9, possibly vacuoles that could be a sign of senescence [58]; Scale = 2 µm. **(B)** Volumes of individual plastids (green, right axis) and their pyrenoid (magenta, left axis) within algal prey (n = 12 plastids), fed host (n = 14 plastids from three hosts) and a 9-week starved host (n = 14 plastids from two hosts). This shows a significant decrease of the plastid volume over time but the pyrenoid volume tended to remain relatively stable. Note that these 9-week-old plastids were still larger than the ones in the algal prey. **(C)** Chlorophyll per plastid volume (ng Chl Plastid-Volume^-1^) was assessed from *P. antarctica* (prey), stolen states in fed and starved (9 weeks) hosts (Chlorophyll measured on triplicate cultures). **(D)** NanoSIMS ^13^C/^12^C isotope ratio image of starved host cell (week 9), showing ^13^C enrichment (log) in several host structures, so photosynthetically-fixed carbon was transferred to the host cell. Stolen plastids (kp) and their pyrenoids (kpy) exhibited lower ^13^C enrichment similar to the host nucleus (hn). Highly ^13^C labeled unknown vacuoles in the symbiosome were observed. The very low content of sulfur and phosphorus in these vacuoles suggests that it could be storage of carbohydrates that are produced by photosynthesis and accumulate over time (Figure S7). Scale = 2 µm. **(E)** EM slice of a 9-week starved host overlayed with 3D reconstruction of stolen plastids (green), pyrenoids (magenta) and thread-like network of stolen mitochondria (red); host nucleus (hn); Scale = 2 µm. **(F)** Starved host cell (week 9) with one stolen plastid surrounded by stolen algal mitochondria (scale = 2 µm) and a zoomed-in view with spherical vacuoles as in (A) (yellow) (Scale = 1µm). See also Tables S1 and S2.

### Carbon fixation by stolen plastids during starvation

The gradual decrease of photosynthetic activity of stolen plastids during starvation raises the question about whether they are still able to fix and translocate carbon to the host. Contrary to the thylakoids and stroma, the pyrenoid volume was comparable between week 9 (1.20 µm^3^ ± 0.44, n= 14) and freshly engulfed plastids (1.19 µm^3^ ± 0.45, n= 14). As a consequence, pyrenoid occupancy in plastids was higher (17% *vs* 11% in fed hosts) (Table S1). Western blot analyses showed that RbcL was still present in stolen plastids at weeks 3 and 9, suggesting the ability to maintain carbon fixation (Figure S2). NanoSIMS ^13^C mapping confirmed that carbon fixation in plastids (low enrichment levels; ∼1500‰), as well as carbon transfer to the host (including nucleus and trichocysts) still took place after 9 weeks of starvation (Figure 5D). Of note, high ^13^C enrichment (∼5000-9000‰) was observed in some vacuoles, localized in the symbiosome, suggesting carbon storage that may accumulate over time (Figures 5D and S6). Very low content of ^32^S, ^31^P and ^12^C^14^N as unveiled by NanoSIMS elemental images was found in these vacuoles, supporting this hypothesis of carbon storage (Figure S7). Thus, despite decreased photosynthetic activity and volume during starvation, stolen plastids were still able to fix and transfer carbon to the host, likely providing nutritional benefits after several months.

Except for the two plastid-encoded Rubisco subunits, all Calvin–Benson–Bassham (CBB) genes are encoded in the nucleus, raising the question of how carbon fixation still takes place. Transcriptomic analyses identified plastidial isoforms of several CBB transcripts (Table S3), all of dinoflagellate origin. The N-terminal extensions of most of these CBB transcripts contain the hallmarks of peridinin-plastid targeting peptides [36], suggesting targeting to the remnant/cryptic peridinin plastid, although such a relict plastid has not been identified in RSD. However, as the NanoSIMS analysis confirmed continued carbon fixation, we consider here the possibility of dual targeting to the peridinin as well as stolen plastids, which presumably co-exist in the same host cell.

### Retention of the algal mitochondrion for several months

Remarkably, the algal mitochondrion was still observed inside host cells after several months of starvation. At week 9, the mitochondrion formed a thread-like network connecting different stolen plastids (Figure 5E) or was closely associated with a single remaining plastid (Figure 5F). At week 30, the mitochondria were still associated with stolen plastids with normal cristae (Figure S6). We hypothesize that the retention of the algal mitochondrion and its intimate interaction with the plastid may contribute to the longevity and functioning of plastids (e.g., ATP and NADPH supply, metabolite exchange, ROS balance, CO_2_ delivery) [37, 38]. To our knowledge, this observation revealing long-term retention of an algal mitochondrion without its original nucleus is unprecedented and unveils a new strategy in kleptoplastidy [39]. Like for the plastid, the transcriptome of the fed host was analyzed to assess the genetic basis of mitochondrion maintenance. We identified all but one protein of the nuclear-encoded Tricarboxylic acid cycle (Krebs cycle; TCA), but all were of dinoflagellate origin. This suggests that, like for the CBB cycle, no gene transfers from the current kleptoplast or former associations have occurred in this pathway. However, with the exception of two candidates, more than one isoform of each TCA protein was identified, which allows for the possibility of dual targeting: “alga-mitochondria targeted” and “host-mitochondria targeted” isoforms (Table 4).

### Worldwide geography of the dinoflagellate host in polar regions

The results of our multimodal study clearly demonstrate the important impact of kleptoplastidy on carbon fluxes at the cellular scale in this association. In order to determine the potential ecological significance of this strategy, originally found in the Ross Sea (Antarctica), we searched for the 18S rRNA V4 sequence marker of RSD in the EukBank database [41] that contains metabarcoding samples from worldwide oceanographic expeditions. This analysis showed that the RSD dinoflagellate is not only found in different regions of the Southern Ocean, but is also widespread in the Arctic (Figure 6). We carried out similar analyses for *Phaeocystis antarctica* and *Phaeocystis pouchetii* (the endemic *Phaeocystis* species in the Antarctic and Arctic regions respectively), which revealed their presence in most samples where RSD sequences were detected during the polar spring and summer. We hypothesize that the RSD dinoflagellate is present in polar regions, including the Arctic, relying on the abundant polar *Phaeocystis* species as prey. These results further highlight the significance of this strategy in the ocean plankton. We hypothesize that this (and potentially other) kleptoplastic association(s) play an important ecosystem-level role in primary productivity and carbon cycling in polar regions.

**Figure 6:**
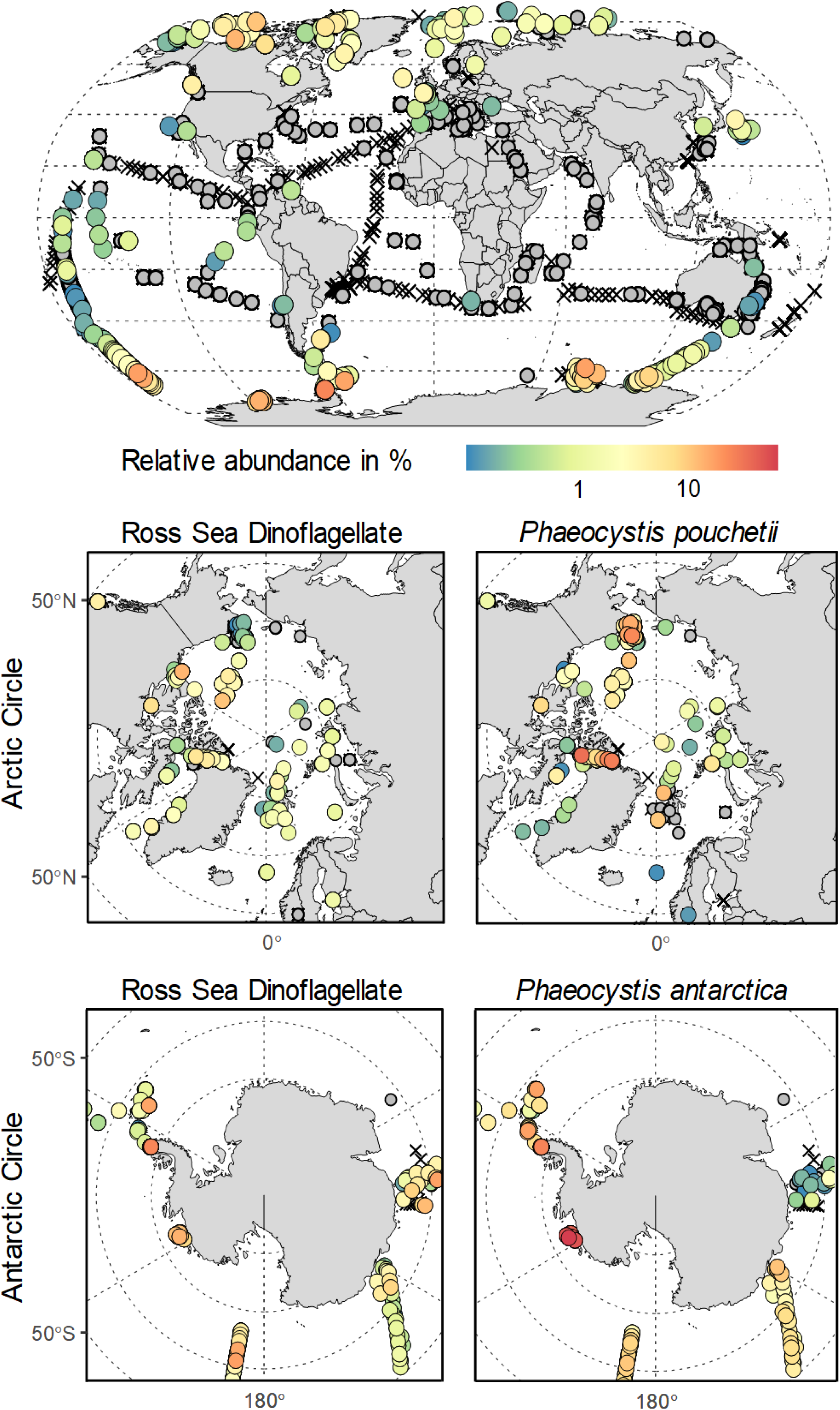
Distribution of kleptoplastidic dinoflagellate RSD as unveiled by environmental metabarcoding datasets. World map showing the relative abundance of metabarcode V4 rDNA reads that matched 100% to query sequences of the RSD dinoflagellate, *Phaeocystis pouchetii* and *Phaeocystis antarctica* (% RSD reads among total reads of the community including protists and metazoans in a given sample) in different stations of the ocean. Higher relative abundance of RSD and polar *Phaeocystis* species reads was observed in the Arctic and Antarctic regions. Note that RSD dinoflagellate was not only found in different regions of Antarctic, but also widely distributed in the Arctic. Grey crosses on the map indicate when RSD or *Phaeocystis* reads were not detected.

## CONCLUSIONS

The combination of photophysiology, 3D electron microscopy and nanoscale carbon flux imaging in this study illuminates, at pertinent temporal and subcellular scales, a new interaction strategy in the plankton whereby the host captures entire algal cells and hijacks expanded nucleus, mitochondrion and plastids. The 6-fold expansion of plastid and C-fixing pyrenoid volumes and boosted photosynthesis, likely supported by the stolen algal nucleus and mitochondrion, is clearly beneficial for the host as we showed that they provide carbon energy. Increase in the volume of the photosynthetic machinery in a foreign host was also found in kleptoplastidic (*Mesodinium rubrum* [17] and *Nusuttodinium aeruginosum* [18]) and photosymbiotic [29] interactions, suggesting the existence of unknown cellular/signaling mechanisms for plastid remodeling. The algal mitochondria also developed into an extensive network in close interaction with plastids and likely contributed to the control and activity of photosynthesis, as shown in diatoms and plants [31–32]. Thus, when algal prey is available, this interaction is more cytoklepty (cell stealing) than kleptoplastidy, whereby a host steals and manipulates most of the organelles of an ecologically successful microalgal cell, such as *Phaeocystis*.

When algal prey is no longer available, however, our study showed that host cells lose the algal nucleus after ∼1 week. Despite the absence of essential algal nucleus-encoded proteins, the stolen plastids and mitochondria remain active for the next few months. This may be explained by the complex genomic toolkit of the host, composed of photosynthesis-related genes inherited i) from the vestigial permanent plastids (peridinin and haptophyte origins), by endosymbiotic gene transfer, and ii) from various algal lineages (including haptophytes) by horizontal gene transfer. Although they still fix and transfer carbon to the host, plastids decrease in volume with diminished photosynthetic activity over several weeks in the absence of the algal nucleus. It is possible that stolen plastids switch to electron cyclic flow mode when PSII is no longer functional.

The host must therefore eventually acquire new plastids from the environment in order to resume vigorous growth. It is clearly advantageous for hosts to rely on an abundant bloom-forming phytoplankton taxon as a source of new organelles, such as *Phaeocystis* (up to 20 million of cells l^-1^) that are often present across seasons [43] [44]. *Phaeocystis* can be present during the austral spring, but reaches extremely high abundances in austral summer (December and January), providing unlimited resources for the host dinoflagellate. During the polar night, the retention time of functional plastids may be extended since degradation could be slower due to the absence of light. In early spring, retained plastids could enable the host to persist until levels of prey are high enough to be encountered and incorporated.

The use of a combination of multiscale approaches with relevant temporal resolution is fundamental for deciphering highly complex cellular interactions in protists and unveiling remarkable mechanisms along the spectrum of plastid endosymbiosis. From an ecological perspective, this study shows the capacity of hosts to hijack plastids and temporarily boost photosynthesis, a process which likely impacts the oceanic carbon cycle. From an evolutionary perspective, the interaction between RSD and *Phaeocystis* provides key insights into cellular and metabolic processes that potentially led to the permanent incorporation of photosynthetic machinery and thus improves our understanding of how eukaryotes can acquire new organelles. This unprecedented example of extended retention of the algal mitochondrion suggests that this organelle may have played a key role in the early stages of plastid acquisition in addition to fundamental endosymbiotic steps (e.g. gene transfer, protein targeting).

## MATERIALS AND METHODS

### Culture conditions of the microalga *Phaeocystis antarctica* and the kleptoplastidy system

A co-culture of *Phaeocystis antarctica* and Ross Sea Dinoflagellates (RSD) isolated from the Ross Sea, Antarctica was maintained in L1 medium [50] (36 ppt salinity) at 4°C with a 12-12 hour light/dark cycle at 45-50 μmol/m^2^/s photon flux intensity. In parallel, the microalga *Phaeocystis antarctica* (called here the prey) was isolated from this co-culture and grown separately in the same conditions. All microscopy images and physiological measurements were performed on solitary cells for accurate counting of cells. Prey *P. antarctica* cells in the exponential phase (∼ day 5) were used to feed RSD cultures. For measurements on “Fed Hosts”, co-cultures were filtered with a 70 µm Corning® Cell strainer, followed by sequential filtrations on a 20 µm mesh fabric to remove prey, RSD clumps and solitary prey cells. Measurements on algal prey were performed on cells filtered with a 40 µm strainer to remove colonies.

For the starvation experiment, the prey-host co-culture was first gently filtered using a 70 µm cell strainer to remove colonies of *P. antarctica,* followed by filtrations on a 20 µm mesh fabric to remove solitary *Phaeocystis* cells and recover RSD dinoflagellate cells (larger than 20 µm in size). These RSD cells were washed three times with cold filtered seawater and once with L1 medium, and transferred to a culture flask. After these filtrations, the presence of *Phaeocystis* (both solitary and colonial stages) was visually checked through an inverted light microscope using both bright field and chlorophyll fluorescence. RSD cells were then placed at 4°C and the same conditions described above (45-50 µmol/m^2^/s with a 12h-12h light-dark cycle). The culture medium was refreshed every 4 weeks during starvation by filtering the host cells on a clean (sonicated) 10 µm mesh fabric followed by washing with cold fresh L1 medium. All subsequent measurements on “Starved Hosts” were performed with no further filtering.

### Photosynthesis measurements and oxygen production

The rapid light curves to measure Electron Transport Rate through PSII (rETR II) were performed using Pulse-Amplitude Modulation (PAM) fluorometry on the microalgae culture *P. antarctica* and the host RSD (fed and starved conditions) with 300 milliseconds saturating pulse followed by 20 seconds between 8 increasing actinic light exposures (15 to 219 µmol photons m^-2^ s^-1^). 500 ml of each sample (with biological triplicates) was pipetted directly from the growth light condition into a WALZ KS-2500 water-jacketed chamber equipped with a stir bar maintained at 4°C by a Mini-PAM-II fluorometer. The instrument is operated with WinControl-3 software (Heinz Walz GmbH). Curve-fitting for ETR values and extraction of the parameters alpha, beta, ETRmax, Ek and ps from the fitted curves was performed using the R package phytotools [51] and to extract the coefficients/parameters alpha, beta, ETRmax, Ek and ps from the fitted curve. A modified Chi-square test was performed on the fitted-curved to compare the curves using the Excel sheet (N>1, mean and SE) from [27]. The first Y(II) value from the rapid light curve was considered for maximum potential quantum efficiency of Photosystem II (Fv/Fm). The samples were dark-adapted for one minute before starting the light curve. Dark-adapting the cells separately for 40 minutes before Fv/Fm measurement did not significantly change the value of Fv/Fm (±0.01 compared to 1-minute dark prior to rapid light curve). This is possibly due to the light level (45-50 μmol/m^2^/s photon flux intensity). For the oxygen measurements, 500 ml of samples was pipetted in a WALZ KS-2500 water-jacketed chamber (Heinz Walz GmbH) paired with a FSO2-1 oxygen meter and optical microsensor - FireStingO2 (PyroScience GmbH) controlled by the WinControl-3 software. Net Oxygen production was measured by exposing cultures to actinic light set at 90 µmol/m^2^/s for 9 minutes followed by 2 minutes in the dark to measure respiration. Gross maximum production was calculated by: O2 Gross = O2 Net – (Respiration (negative slope)).

### Sample Preparation for electron microscopy (EM)

The microalga *Phaeocystis antarctica* and host RSD dinoflagellate cultures in exponential phases were centrifuged at 2000 g for two minutes and cryofixed using High Pressure Freezing (HPM100, Leica Microsystems, Austria) at a pressure of 210 MPa at −196°C as in [29]. This was followed by freeze-substitution (EM ASF2, Leica Microsystems, Austria), where vitrified ice was replaced with dried acetone and 2% osmium tetroxide for TEM and nanoSIMS, with 0.5% uranyl acetate added for FIB-SEM. Samples are embedded in EPON-substitute (for TEM) or Araldite (for SEM). More details of the sample preparation protocols are provided in protocol.io [52]. The resin blocks were then stored in dry conditions prior to imaging.

### Transmission and Scanning EM observations

For transmission electron microscopy (TEM), 70 nm thick sections obtained using a diamond knife and a UC7 ultramicrotome (Leica Microsystems, Austria) were collected on 200 mesh formvar-carbon coated grids. Sections were stained with 4% uranyl acetate (10 min) and lead citrate (5 min). Micrographs were obtained using a Tecnai G2 Spirit BioTwin microscope (FEI) operating at 120 kV with an <u>Orius</u> SC1000 CCD camera (Gatan). For scanning electron microscopy (SEM), 200 nm thick sections were transferred to a silicon wafer and observed at low voltage (3 kV, 800 pA) with a Sense BSD backscattered electron detector in a Zeiss GeminiSEM 460.

### FIB-SEM acquisition, segmentation, and morphometric analyses

Focused Ion beam Scanning Electron Microscopy (FIB-SEM) tomography was performed with a Zeiss CrossBeam 550 microscope (Zeiss, Germany) as [66]. The resin block containing the cells was fixed on a stub with silver paste and surface-abraded with a diamond knife in a microtome to obtain a flat and clean surface. Samples were then metallized with 4-8 nm of platinum to avoid charging during observations. Inside the FIB-SEM, a second platinum layer (1–2 μm) was deposited locally on the analyzed area to mitigate possible curtaining artifacts. The sample was then abraded slice by slice with the Ga+ ion beam (typically with a current of 700 pA at 30 kV). Each exposed surface was imaged by SEM at 1.5 kV and with a current of ∼1 nA using the in-column EsB backscatter detector. Similar milling and imaging modes were used for all samples. Automatic focus and astigmatism correction were performed during image acquisition, typically at approximately hourly intervals. For each slice, a thickness of 6 or 8 nm was removed and SEM images were recorded with a corresponding pixel size of 6 to 8 nm in order to obtain an isotropic voxel size. Entire volumes were imaged with 800–3000 frames, depending on the cell type and volume. The first steps of image processing were performed using Fiji software for registration (adapted Stackreg plugin), and noise reduction (3D mean function of the 3D suite plugin). Raw electron microscopy data have been deposited in the Electron Microscopy Public Image Archive (EMPIAR), accession code EMPIAR-XXX.

Two segmentation strategies (automatic with a new algorithm and semi-automatic) were used to reconstruct and generate models of cells and organelles in three dimensions. The overview of plastid and mitochondria morphologies was obtained using the FeatureForest, a napari plugin that uses the Segment Anything Model (SAM) [62] to extract features from images and train a random forest classifier based on these features and inputs from the user. Once trained on a small subset of the data, the random forest model, together with SAM, allows segmenting the entire stack. In order to obtain smoother object segmentation, FeatureForest applies post-processing steps by filling holes in the segmentation, removing connected components smaller than a certain threshold and using the remaining connected components to generate bounding boxes used as input prompts to SAM, yielding well-defined object boundaries. The plugin was created in collaboration with the Jug lab and the Bioimage Analysis Unit of the National Facility for Data Handling and Analysis at the Human Technopole, Milan, Italy through the AI4Life project (https://github.com/ai4life-opencalls/oc_1_project_52). The plugin repository is publicly available at https://github.com/juglab/featureforest/blob/main/docs/index.md.

Semi-automatic and manual segmentation using 3D slicer [48] was used to obtain volumes of organelles. Plastids, pyrenoids, mitochondria and nuclei of the microalga (native and stolen states) and RSD dinoflagellate host cells were segmented to reconstruct the cellular organization of kleptoplastidy and quantify volumes (µm^3^). Each segment (e.g. pyrenoid, plastid, mitochondria) was colored using paint tools, adjusting the threshold range of the pixel values (intensity) of the images. Different views and slices were generated for the model using the EasyClip module. Volume measurements were then calculated using the Segmentation Statistics module. Paraview was also used to edit the 3D models and generate images and videos [49].

### NanoSIMS experiment with labeled ^13^C-bicarbonate: correlative imaging and analysis

NanoSIMS experiments were conducted on two culture conditions: fed and starved RSD dinoflagellate host cells. For the fed conditions, around 1 ml of prey microalgae (*P. antarctica* in exponential phase) was added one week before cryofixation to culture flasks containing 90 ml RSD culture. For the starved conditions, RSD dinoflagellates were washed several times to remove the *Phaeocystis* prey as explained above and maintained for 9 weeks without prey before cryofixation for nanoSIMS. For both conditions, cells were incubated with stable isotopes for 24 hours. More specifically, 10 ml of an isotopic solution with 10X concentration containing H^13^CO_3_ was added to the culture flasks. A control culture without isotope labeling was also maintained in the same culture conditions. After 24h incubation with H^13^CO_3_, cells were harvested by centrifugation for two minutes at 2000 g and cryofixed with High Pressure Freezing (see above). Thin sections (100 nm - 200 nm) were first imaged using TEM or SEM in order to observe the ultrastructure of the RSD dinoflagellates containing stolen plastids and identify the cells of interest for isotopic analysis. The TEM grids or SEM silicon wafers were then introduced into a NanoSIMS ion microprobe (Cameca, Gennevilliers, France) at either EPFL, Lausanne (Switzerland) or at Institut Curie, Orsay (France) and analyses were performed on the preselected cells. The instrument delivers a primary beam of Cesium (Cs+) ions to the sample surface, focused to a ∼50-100nm spot. The secondary ions are extracted from the sample surface and transferred to a high-resolution mass-spectrometer.

For NanoSIMS on fed host cells, the secondary ^12^C^14^N^-^ and ^13^C^14^N^-^ ions were collected in electron multipliers at the exit of the mass spectrometer with a mass resolution (M/ΔM) of ∼ 9000. Each capture consisted of six sequential images. For 9-week starved cells, five ion species (^12^C_2_^-^, ^12^C^13^C^-^, ^12^C^14^N^-^, ^31^P^-^ and ^32^S^-^) were collected in parallel, also captured in multiframe acquisition mode. Individual frames were recorded at 512 x 512 pixel definition, with a dwell time of 0.5 ms per pixel, with up to 100 frames accumulated for each acquisition. The collection of C_2_^-^ was preferred to C^-^ for enhanced ion emission yield thus higher count rate and improved counting statistics for ^13^C quantification. However, appropriate mass resolving power has to be applied to ensure specific detection of ^12^C^13^C^-^ ions against the interference ^12^C_2_ ^1^H^-^ ions (although at a moderate relative mass separation, M/ΔM, of 5600, but with an intensity of 10 times that of ^12^C^13^C^-^). Sequential images acquired from the NanoSIMS instruments were aligned frame-to-frame with the L’IMAGE software (developed by Larry Nittler; marketed by the Carnegie Institution of Washington, USA) for drift compensation. The ratio images for the carbon isotope were then obtained by pixel-to-pixel division of accumulated ^12^C^15^N^-^ and ^12^C^14^N^-^, or ^12^C^13^C^-^ and ^12^C_2_ ^-^ images. The resulting isotopic images were then superimposed on the TEM/SEM images to precisely measure the isotopic ratios in specific subcellular locations thus allowing us to study carbon allocation within the RSD [17].

### Ultrastructural expansion microscopy (U-ExM) and Confocal Microscopy

Dinoflagellate RSD cells were gently pelleted at 4°C by centrifugation at 1000 g (3170 rpm) for 3 minutes. The pellets were fixed in a final concentration of 4% formaldehyde for 20 minutes at room temperature. Pellets were then washed twice with 1 ml 1x PBS followed by centrifugation at 1000 g for 3 minutes at 20°C. Finally, pelleted cells were resuspended in 1 ml of 1x PBS and stored at 4°C until expansion. Expansion was performed as previously described [63–65]. Fixed cells were either attached to a poly-D-Lysine coverslip or sedimented in a microtube before being cross-linked in AA/FA solution (1% acrylamide (AA)/0.7% formaldehyde (FA)) for 12 hours at 37°C. The cells in microtubes were then allowed to sediment to the bottom of a 12-well plate, and excess liquid was carefully removed with minimal disturbance. Gelation was performed using a monomer solution consisting of 19% (wt/wt) sodium acrylate (Combi-Blocks, ref: QC-1489), 10% (wt/wt) acrylamide (Sigma-Aldrich; ref: A4058), and 0.1% (wt/wt) N, N&#39;-methylenebisacrylamide (Sigma-Aldrich; ref: M1533) in PBS. The process was carried out at 37°C for 1 hour in a moist chamber. Gel denaturation was performed in denaturation buffer (50 mM Tris pH 9.0, 200 mM NaCl, 200 mM SDS, pH to 9.0) for 15 min at RT. The gel was then incubated at 95°C for 1.5 hours. Following denaturation, expansion was performed with several water rinses as previously described [63–65]. Post expansion, gel diameter was measured and used to determine the expansion factor (4.1875X). Scale bars in the images are corrected to indicate actual size. Sequential antibody staining was performed for imaging thylakoids with pyrenoids and protein localization (Figure 1C). Antibodies were prepared in PBS supplemented with 3% BSA and 0.1% Tween20. For thylakoid staining, a rabbit primary antibody targeting Photosystem II PsbA (Ref: AS05 084, Agrisera) was used at a 1/250 dilution. The primary antibody was tagged with an Invitrogen Donkey-anti-Rabbit IgG (H+L) Highly Cross-Adsorbed Secondary Antibody, Alexa Fluor™ 488 (Ref: A-21206, Thermo Fisher) at a 1/250 dilution. The secondary antibody was washed 5 times with PBS supplemented with 3% BSA and 0.1% Tween20. Subsequently, pyrenoid staining was performed using a rabbit primary antibody targeting RuBisCO (Ref: AS03 037, Agrisera) at a 1/250 dilution. This primary antibody was tagged with a different Invitrogen Donkey-anti-Rabbit IgG (H+L) Highly Cross-Adsorbed Secondary Antibody, Alexa Fluor™ 568 (Ref: A10042, Thermo Fisher) at 1/500 dilution. For other protein localization (Figure S1), primary antibodies (beta-tubulin antibody, Ref: AA344-R, Geneva Antibody Facility. Alpha-tubulin antibody, Ref: AA345-R, Geneva Antibody Facility. Centrin2 (20H5) antibody, Ref: 04-1624, Merck. RuBisCO and Photosystem II PsbA antibodies, see above) were incubated overnight at 37°C at dilutions of 1:250 to 1:500. After three washes in 0.1% PBST, secondary antibodies (Invitrogen Donkey or Goat-anti-Rabbit and anti-Mouse IgG (H+L) Highly Cross-Adsorbed Secondary Antibody, Alexa Fluor™ 488 or 647 Plus Ref: A21206, A28175, A32795, A32728, Thermo Fisher) were added at 1:500 to 1:1000 and incubated for 4hrs at 37°C. For protein pan-labelling, gels were incubated with Alexa Fluor™ 405 or 594 NHS Ester (Ref: A30000 and A20004, Thermofisher) at 1/500 dilution in NaHCO3 (pH=8.28) for 1.5 hrs. For staining of nuclei of the dinoflagellate and algal prey, BioTracker NIR694 Nuclear Dye (Ref: SCT118, Merck Millipore) or Hoechst 33342 (Ref: 62249, Thermo Fisher) was incubated at a dilution of 1/500 and 1/1000, respectively, in PBS for 30 minutes before re-expansion in water.

For gel mounting, gels were cut to appropriate sizes and attached to pre-coated Poly-D-lysine coverslips and sealed using i-Spacers (Ref: #IS013, Sunjin Lab). For imaging of expanded samples (Figures 1C and Figure S1), an upright Leica SP8 confocal microscope with an HC PL APO 40X/1.25 Glycerol objective was used. For confocal microscopy of non-expanded samples, cells were allowed to adhere to a 35 mm Poly-D-lysine-coated glass-bottom Petri dish for 15 minutes at 4°C. Z-stack images of cells were imaged with a Zeiss LSM880 inverted confocal microscope equipped with a Zeiss Plan-Apochromat 963 (1.4) Oil DIC M27 objective, and Zeiss Airyscan super-resolution detector.

### Transcriptomic analysis and prediction of signal peptide origin

Known CBB cycle proteins from several eukaryotic model organisms were selected from NCBI, while myzozoan representatives from the TCA cycle were obtained from (https://bmcbiol.biomedcentral.com/articles/10.1186/s12915-021-01007-2). Candidates were used as BLASTP queries against a comprehensive custom protein database containing representatives from most major eukaryotic groups, with a focus on plastid-containing lineages plus selected taxa from non-plastidial lineages and RefSeq data from all bacterial phyla at NCBI (last accessed December 2017). The database was subjected to CD-HIT with a similarity threshold of 85% to reduce redundant sequences and paralogs, except for the two RSD data sets created in [22] that were clustered at 98%. Of note, as described in [22], the RSD transcriptome is substantially contaminated by the prey *Phaeocystis antarctica*. A cleaned dataset, ‘RSD allclean noPhaeo’ was therefore generated and used in conjunction with the uncleaned dataset ‘RSD Temp01’. Search results of the BLASTP step were parsed for hits with an e-value threshold ≤1e-25 and a query coverage of ≥ 50% to reduce the possibility of paralogs and short sequences. The number of bacterial hits was restrained to 20 hits per phylum (for FCB group, most classes of Proteobacteria, PVC group, Spirochaetes, Actinobacteria, Cyanobacteria (unranked) and Firmicutes) or 10 per phylum (remaining bacterial phyla) as defined by NCBI taxonomy.

Parsed hits of queries corresponding to the same protein were combined, deduplicated and aligned with MAFFT, using the --auto and the --reorder option (https://academic.oup.com/mbe/article/30/4/772/1073398), phylogenetically informative sites were identified with ClipKIT, using default options (10.1371/journal.pbio.3001007) and initial Maximum likelihood tree reconstructions were performed with FastTree v. 2.1.7, using default options (https://www.ncbi.nlm.nih.gov/pmc/articles/PMC2835736/). Due to the complexity of CBB cycle protein phylogenies, the BLASTP step against the custom database was repeated with the initial query protein set plus all RSD orthologs identified in the initial phylogeny, to ensure a comprehensive number of hits for sequences highly similar to RSD. Resulting phylogenies and underlying alignments were inspected manually to remove contaminations and poor-quality sequences in several iterations, repeating the tree reconstruction as described above. Cleaned alignments were then aligned with MAFFT L-INS-i, using the --reorder option, before trimming again with ClipKIT, using default options. Final trees were calculated with IQ-TREE v. 1.6.12 (https://doi.org/10.1093/molbev/msu300), using the LG model, with branch support assessed with 1000 ultrafast bootstrap replicates (https://doi.org/10.1093/molbev/msx281). Bootstraps for the TPI phylogeny did not converge even after increasing to 10,000 iterations. All phylogenies have been deposited to a repository as pdf figures and in newick format (preliminary links: https://figshare.com/s/91c1a6c484d7e03fa61e for the TCA cycle phylogenies and https://figshare.com/s/7d005e093646410a6fb9 for the CBB cycle phylogenies).

To investigate N-terminal extensions and thus intracellular location of RSD sequences, dinoflagellate and haptophyte clades containing RSD sequences were isolated from the corresponding alignments, realigned using MAFFT L-INS-I and then manually inspected for completeness of the sequences as well as for similarity in the N-terminal region to known plastid-targeted sequences of peridinin and haptophyte plastid-harboring dinoflagellates. Prediction of signal peptides as part of N-terminal bipartite leader sequences was performed with SignalP-6.0 selecting “Eukarya” as organism and “Slow” as the model mode (https://pubmed.ncbi.nlm.nih.gov/34980915/). Mitochondrial transit peptides and haptophyte plastid transit peptides were predicted with TargetP-2.0 (doi:10.26508/lsa.201900429). To predict N-terminal transmembrane domains, typical for peridinin-plastid transit peptides, DeepTMHMM (https://www.biorxiv.org/content/10.1101/2022.04.08.487609v1) and Phobius, as implemented in InterProScan (https://europepmc.org/article/MED/24451626), were used. Curated alignments underlying the plastidial clades of CBB cycle proteins containing RSD representatives have been deposited to a repository (preliminary link: https://figshare.com/s/3bce42bce704e90ec5a6).

### Protein extraction and Western Blot

Total protein extracts were obtained from prey *Phaeocystis antarctica*, and fed and starved RSD dinoflagellate hosts at weeks 1, 3 and 9 in 50 mM Tris buffer pH 8.0 supplemented with protein inhibitor cocktail (539131, Calbiochem). A Precellys device (Bertin Technologies) and micro glass beads (500 µm) were used to disrupt cells with two 30 second cycles at 5000 rpm. After centrifugation, proteins in the supernatant were precipitated overnight at −20°C in 100 % acetone. After a second centrifugation, the pellet was solubilized for 5 min (RT) in 50 mM Tris (pH 6.8), 2% sodium dodecyl sulphate, 10 mM EDTA, and protein inhibitor cocktail. After a second centrifugation, supernatant was retained and protein quantified with the DC Protein assay kit II (Biorad). Protein samples (3µg - 5µg) were loaded on 10% SDS-PAGE gels (Mini-PROTEAN TGX Precast Protein Gels, Biorad) and blotted onto nitrocellulose membranes. Membranes were blocked for 1h with 5% low fat milk powder in TBS-T Tween 0.1% and probed with antiRBCL antibody (AS03 037), antiPsbA (PSII - chloroplast encoded subunit, AS05 084), antiPsaC (PSI - chloroplast encoded,AS10 939) (for Gels 1 and 2), antiPsbO (PSII - nuclear encoded subunit, AS21 4689) (dilution for all mentioned above: 1:10000 TBS-T, overnight) and antiH3 antibody (Histone 3, internal control AS10 710, dilution 1/5000 in TBS-T). For the secondary antibody, HRP conjugated anti rabbit antibody (111-035-003) (Interchim, 1:10000, 1h) in TBS-T was used. Antibody incubations were followed by washing in TBS-T. All steps were performed at room temperature with agitation. Blots were developed for 1min with the ECL Prime detection kit (RPN2232, Amersham) according to the manufacturer’s instructions (GE Heathcare). Images of the blot were obtained using a CCD imager (Chemidoc MP system, Biorad) and ImageJ software.

### 18S rDNA sequencing and global biogeographic analyses using metabarcoding datasets

Cultures of the RSD without *Phaeocystis* prey were centrifuged at 7500 rpm for 5 minutes and the supernatant was gently removed with a pipette and filtered tips. Then, 20µl of the buffer Phire of the Plant Direct PCR Master Mix (Thermofisher) was added and the DNA sample was incubated at 95°C for 5 minutes. PCR amplification and sequencing of the SSU region was performed using primers as described in [14]. To obtain the entire 18S rDNA sequence of the RSD, three different primer sets were used with the Phire Plant Direct PCR Master Mix (Thermofisher) in the following conditions: 30s et 98°C pre-denaturation, followed by 35 cycles of 10s at 98°C, 30s at 55°C and 30s at 72°C, with a final extension of 10 min at 72°C. PCR products were then purified using the QIAquick gel extraction kit (QIAGEN) after electrophoresis on 1% agarose gel. Sequences were manually aligned and edited using SnapGene software and were deposited in GenBank as a single contig of 1784 bp (GenBank accession no. = XXXX).

Prior to metabarcoding analyses, we verified that the 18S rDNA sequence of RSD dinoflagellate has a unique V4 region sequence among the family Kareniaceae, including its closest relative *Shimiella gracilenta* (a kleptoplastidic dinoflagellate with a cryptophyte as prey [14]). Global biogeographic analysis of the dinoflagellate host RSD was carried out using the EukBank metabarcoding dataset of the V4 hypervariable region of the 18S rRNA accessed at doi: 10.5281/zenodo.7804946. The V4 sequence of the RSD host was used to search reads with 100% match in the EukBank dataset [41] and relative abundance of reads (number of reads from RSD divided by the total number of reads in the sample) was assessed. The same approach was conducted with the V4 region of the 18S rDNA sequences of *Phaeocystis antarctica* and *Phaeocystis pouchetii* retrieved on GenBank (KF925339 for *P. antarctica* and AF182114 for *P. pouchetii*) to investigate their global biogeography in the EukBank dataset.

## ACKNOWLEDGMENTS

Research was supported by the ANR EPHEMER and the ERC consolidator grant SymbiOCEAN (101088661). We acknowledge CurieCoreTech, the scientific and technological platforms within Institut Curie, for the use of Curie-NanoSIMS as well as the NanoSIMS platform in Lausanne (Anders Meibom and Stephane Escrig). FIB-SEM and SEM imaging were carried out on the Platform for Nanocharacterisation (PFNC) of CEA-Grenoble, supported by the “Recherche Technologique de Base” and “France 2030 - ANR-22-PEEL-0014” programs of the French National Research Agency (ANR). We thank Dr Christine Moriscot from the IBS/ISBG EM platform, part of the Grenoble Instruct-ERIC center (ISBG; UMS 3518 CNRS-CEA-UGAEMBL) within the Grenoble Partnership for Structural Biology (PSB), supported by FRISBI (ANR-10-INBS-05-02), and GRAL, financed within the University Grenoble Alpes graduate school (Ecoles Universitaires de Recherche) CBH-EUR-GS (ANR-17-EURE-0003). The IBS/ISBG EM facility is led by Dr Guy Schoehn and supported by the AuvergneRhône-Alpes Region, the Fondation Recherche Medicale (FRM), the fonds FEDER and the GIS-Infrastructures en Biologie Sante et Agronomie (IBISA). MS and the AI4Life Horizon Europe Program Consortium received funding from the European Commission through the Horizon Europe Program (AI4LIFE project with grant agreement 101057970-AI4LIFE). We are grateful to Nicolas Henry and the Roscoff Bioinformatics platform ABiMS (http://abims.sb-roscoff.fr), part of the Institut Français de Bioinformatique (ANR-11-INBS-0013) and BioGenouest network, for providing help and/or computing and/or storage resources. EH is funded by and Lumina Quaeruntur Grant of the Czech Academy of Sciences (LQ200962204). OD and MO are core funded by the Department of Biochemistry at University of Geneva and an SNSF Starting Grant 2023 (TMSGI3_218007). We thank Ian Probert for his feedback on the manuscript.

## AUTHOR CONTRIBUTIONS

A.K.R. and J.D. designed research and write the manuscript. B.G., J.D. and A.K.R. jointly performed the sample preparation for electron microscopy. C.L. and T.W. conducted nanoSIMS experiments, and T.W. and A.K.R. processed and interpreted data. A.K.R. and D.Y. carried out and interpreted the photophysiology measurements. M.P. conducted rDNA sequencing. M.O. and O.D. performed the expansion microscopy and E.H. the transcriptomic analyses. PH.J acquired and pre-processed the FIB-SEM imaging data. M.S., A.K.R. and J.Deschamps. contributed to the image analysis from FIB-SEM data. A.K.R. and F.C. performed the protein extraction and western blots analyses. R.G. helped to draft the manuscript.

## REFERENCES

[1] Douglas, A. E. (2014). Symbiosis as a general principle in eukaryotic evolution. Cold Spring Harbor Perspectives in Biology, 6(2), a016113.

[2] Worden, A. Z., Follows, M. J., Giovannoni, S. J., Wilken, S., Zimmerman, A. E., & Keeling, P. J. (2015). Rethinking the marine carbon cycle: factoring in the multifarious lifestyles of microbes. Science, 347(6223), 1257594.

[3] Archibald JM. The puzzle of plastid evolution. Curr Biol. 2009 Jan 27;19(2):R81–8. doi: 10.1016/j.cub.2008.11.067. PMID: 19174147.

[4] Keeling, P. J. (2014). The impact of history on our perception of evolutionary events: endosymbiosis and the origin of eukaryotic complexity. Cold Spring Harbor perspectives in biology, 6(2), a016196.

[5] Christa, G., Zimorski, V., Woehle, C., Tielens, A. G., Wägele, H., Martin, W. F., & Gould, S. B. (2014). Plastid-bearing sea slugs fix CO2 in the light but do not require photosynthesis to survive. Proceedings of the Royal Society B: Biological Sciences, 281(1774), 20132493.

[6] Van Steenkiste, N. W., Stephenson, I., Herranz, M., Husnik, F., Keeling, P. J., & Leander, B. S. (2019). A new case of kleptoplasty in animals: marine flatworms steal functional plastids from diatoms. Science Advances, 5(7), eaaw4337.

[7] Maselli, M., Anestis, K., Klemm, K., Hansen, P. J., & John, U. (2021). Retention of prey genetic material by the kleptoplastidic ciliate *Strombidium cf. basimorphum*. Frontiers in Microbiology, 12, 694508.

[8] Nishitani, G., Nagai, S., Hayakawa, S., Kosaka, Y., Sakurada, K., Kamiyama, T., & Gojobori, T. (2012). Multiple plastids collected by the dinoflagellate *Dinophysis mitra* through kleptoplastidy. Applied and Environmental Microbiology, 78(3), 813–821.

[9] Gast, R. J., Moran, D. M., Dennett, M. R., & Caron, D. A. (2007). Kleptoplasty in an Antarctic dinoflagellate: caught in evolutionary transition?. Environmental Microbiology, 9(1), 39–45.

[10] Karnkowska, A., Yubuki, N., Maruyama, M., Yamaguchi, A., Kashiyama, Y., Suzaki, T., … & Leander, B. S. (2023). Euglenozoan kleptoplasty illuminates the early evolution of photoendosymbiosis. Proceedings of the National Academy of Sciences, 120(12), e2220100120.

[11] Pinko, D., Abramovich, S., Rahav, E., Belkin, N., Rubin-Blum, M., Kucera, M., … & Abdu, U. (2023). Shared ancestry of algal symbiosis and chloroplast sequestration in foraminifera. Science advances, 9(41), eadi3401.

[12] Sørensen, M. E., Zlatogursky, V. V., Onuţ-Brännström, I., Walraven, A., Foster, R. A., & Burki, F. (2023). A novel kleptoplastidic symbiosis revealed in the marine centrohelid Meringosphaera with evidence of genetic integration. Current Biology, 33(17), 3571–3584.

[13] Chihara, S., Nakamura, T., & Hirose, E. (2020). Seasonality and longevity of the functional chloroplasts retained by the sacoglossan sea slug *Plakobranchus ocellatus* van Hasselt, 1824 inhabiting a subtropical back reef off Okinawa-jima Island, Japan. Zoological Studies, 59.

[14] Ok, J. H., Jeong, H. J., Lee, S. Y., Park, S. A., & Noh, J. H. (2021). *Shimiella* gen. nov. and *Shimiella gracilenta* sp. nov.(Dinophyceae, Kareniaceae), a kleptoplastidic dinoflagellate from Korean waters and its survival under starvation. Journal of Phycology, 57(1), 70–91.

[15] Waller, R. F., & Kořený, L. (2017). Plastid complexity in dinoflagellates: a picture of gains, losses, replacements and revisions. In Advances in botanical research (Vol. 84, pp. 105-143). Academic Press.

[16] Cruz, S., LeKieffre, C., Cartaxana, P., Hubas, C., Thiney, N., Jakobsen, S., … & Meibom, A. (2020). Functional kleptoplasts intermediate incorporation of carbon and nitrogen in cells of the Sacoglossa sea slug *Elysia viridis*. Scientific reports, 10(1), 10548.

[17] Johnson, M. D., Moeller, H. V., Paight, C., Kellogg, R. M., McIlvin, M. R., Saito, M. A., & Lasek-Nesselquist, E. (2023). Functional control and metabolic integration of stolen organelles in a photosynthetic ciliate. Current Biology, 33(5), 973–980.

[18] Onuma, R., & Horiguchi, T. (2015). Kleptochloroplast enlargement, karyoklepty and the distribution of the cryptomonad nucleus in *Nusuttodinium* (= Gymnodinium) *aeruginosum* (Dinophyceae). Protist, 166(2), 177–195.

[19] Yamada, N., Bolton, J. J., Trobajo, R., Mann, D. G., Dąbek, P., Witkowski, A., … & Kroth, P. G. (2019). Discovery of a kleptoplastic ‘dinotom’dinoflagellate and the unique nuclear dynamics of converting kleptoplastids to permanent plastids. Scientific reports, 9(1), 10474.

[20] Garric, S., Ratin, M., Marie, D., Foulon, V., Probert, I., Rodriguez, F., & Six, C. (2024). Impaired photoacclimation in a kleptoplastidic dinoflagellate reveals physiological limits of early stages of endosymbiosis. Current Biology, 34(14), 3064–3076.

[21] Sellers, C. G., Gast, R. J., & Sanders, R. W. (2014). Selective feeding and foreign plastid retention in an Antarctic dinoflagellate. Journal of phycology, 50(6), 1081–1088.

[22] Hehenberger, E., Gast, R. J., & Keeling, P. J. (2019). A kleptoplastidic dinoflagellate and the tipping point between transient and fully integrated plastid endosymbiosis. Proceedings of the National Academy of Sciences, 116(36), 17934–17942.

[23] Larkum, A. W., Lockhart, P. J., & Howe, C. J. (2007). Shopping for plastids. Trends in Plant Science, 12(5), 189–195.

[24] Keeling, P. J. (2013). The number, speed, and impact of plastid endosymbioses in eukaryotic evolution. Annual review of plant biology, 64(1), 583–607.

[25] Stamatakis, K., Vayenos, D., Kotakis, C., Gast, R. J., & Papageorgiou, G. C. (2017). The extraordinary longevity of kleptoplasts derived from the Ross Sea haptophyte *Phaeocystis antarctica* within dinoflagellate host cells relates to the diminished role of the oxygen-evolving Photosystem II and to supplementary light harvesting by mycosporine-like amino acid/s. Biochimica et Biophysica Acta (BBA)-Bioenergetics, 1858(2), 189–195.

[26] Genty, B., Briantais, J. M., & Baker, N. R. (1989). The relationship between the quantum yield of photosynthetic electron transport and quenching of chlorophyll fluorescence. Biochimica et Biophysica Acta (BBA)-General Subjects, 990(1), 87–92.

[27] Hristova, K., & Wimley, W. C. (2023). Determining the statistical significance of the difference between arbitrary curves: A spreadsheet method. Plos one, 18(10), e0289619.

[28] He, S., Crans, V. L., & Jonikas, M. C. (2023). The pyrenoid: the eukaryotic CO2-concentrating organelle. The Plant Cell, 35(9), 3236–3259.

[29] Uwizeye, C., Mars Brisbin, M., Gallet, B., Chevalier, F., LeKieffre, C., Schieber, N. L., … & Decelle, J. (2021). Cytoklepty in the plankton: a host strategy to optimize the bioenergetic machinery of endosymbiotic algae. Proceedings of the National Academy of Sciences, 118(27), e2025252118.

[30] Lütz, C., & Engel, L. (2007). Changes in chloroplast ultrastructure in some high-alpine plants: adaptation to metabolic demands and climate?. Protoplasma, 231, 183–192.

[31] Bailleul, B., Berne, N., Murik, O., Petroutsos, D., Prihoda, J., Tanaka, A., … & Finazzi, G. (2015). Energetic coupling between plastids and mitochondria drives CO2 assimilation in diatoms. Nature, 524(7565), 366–369.

[32] Mueller-Schuessele, S. J., & Michaud, M. (2018). Plastid transient and stable interactions with other cell compartments. Plastids: Methods and Protocols, 87-109.

[33] Boisvert, F. M., Van Koningsbruggen, S., Navascués, J., & Lamond, A. I. (2007). The multifunctional nucleolus. Nature reviews Molecular cell biology, 8(7), 574–585.

[34] Dorrell, R. G., & Howe, C. J. (2015). Integration of plastids with their hosts: Lessons learned from dinoflagellates. Proceedings of the National Academy of Sciences, 112(33), 10247–10254.

[35] Domínguez, F., & Cejudo, F. J. (2021). Chloroplast dismantling in leaf senescence. Journal of Experimental Botany, 72(16), 5905–5918.

[36] Patron, N. J., Waller, R. F., Archibald, J. M., & Keeling, P. J. (2005). Complex protein targeting to dinoflagellate plastids. Journal of Molecular Biology, 348(4), 1015–1024.

[37] Mueller-Schuessele, S. J., & Michaud, M. (2018). Plastid transient and stable interactions with other cell compartments. Plastids: Methods and Protocols, 87-109.

[38] Cardol, P., Alric, J., Girard-Bascou, J., Franck, F., Wollman, F. A., & Finazzi, G. (2009). Impaired respiration discloses the physiological significance of state transitions in Chlamydomonas. Proceedings of the National Academy of Sciences, 106(37), 15979–15984.

[39] Kim, E., & Archibald, J. M. (2010). Plastid evolution: gene transfer and the maintenance of’stolen’organelles. BMC biology, 8, 1–3.

[40] Novák Vanclová, A. M., Nef, C., Füssy, Z., Vancl, A., Liu, F., Bowler, C., & Dorrell, R. G. (2024). New plastids, old proteins: repeated endosymbiotic acquisitions in kareniacean dinoflagellates. EMBO reports, 25(4), 1859–1885.

[41] Berney, C., Mahé, F., Henry, N., Lara, E., de Vargas, C., EukBank consortium, 2023. EukBank 18S V4 dataset [Data set]. Zenodo. doi:10.5281/zenodo.7804946

[42] Johnson, M. D., Oldach, D., Delwiche, C. F., & Stoecker, D. K. (2007). Retention of transcriptionally active cryptophyte nuclei by the ciliate *Myrionecta rubra*. Nature, 445(7126), 426–428.

[43] Smith Jr, W. O., Dennett, M. R., Mathot, S., & Caron, D. A. (2003). The temporal dynamics of the flagellated and colonial stages of *Phaeocystis antarctica* in the Ross Sea. Deep Sea Research Part II: Topical Studies in Oceanography, 50(3-4), 605–617.

[44] Smith Jr, W. O., & Jones, R. M. (2015). Vertical mixing, critical depths, and phytoplankton growth in the Ross Sea. ICES Journal of Marine Science, 72(6), 1952–1960.

[45] Gambarotto, D., Zwettler, F. U., Le Guennec, M., Schmidt-Cernohorska, M., Fortun, D., Borgers, S., … & Guichard, P. (2019). Imaging cellular ultrastructures using expansion microscopy (U-ExM). Nature methods, 16(1), 71–74.

[46] Decelle, J., Veronesi, G., LeKieffre, C., Gallet, B., Chevalier, F., Stryhanyuk, H., … & Musat, N. (2021). Subcellular architecture and metabolic connection in the planktonic photosymbiosis between Collodaria (radiolarians) and their microalgae. Environmental Microbiology, 23(11), 6569–6586.

[47] Schmid, B., Schindelin, J., Cardona, A., Longair, M., & Heisenberg, M. (2010). A high-level 3D visualization API for Java and ImageJ. BMC bioinformatics, 11, 1–7.

[48] Fedorov, A., Beichel, R., Kalpathy-Cramer, J., Finet, J., Fillion-Robin, J. C., Pujol, S., … & Kikinis, R. (2012). 3D Slicer as an image computing platform for the Quantitative Imaging Network. Magnetic resonance imaging, 30(9), 1323–1341.

[49] Ahrens, J., Geveci, B., Law, C., Hansen, C., & Johnson, C. (2005). 36-paraview: An end-user tool for large-data visualization. The visualization handbook, 717, 50038–1.

[50] Guillard, R. R. L., & Hargraves, P. E. (1993). Stichochrysis immobilis is a diatom, not a chrysophyte. Phycologia, 32(3), 234–236. 10.2216/i0031-8884-32-3-234.1

[51] Silsbe, G. M., Malkin, S. Y., & Silsbe, M. G. (2015). Package ‘phytotools’.

[52] Gallet, B., Moriscot, C., Schoehn, G., & Decelle, J. (2024). Cryo-fixation and resin embedding of biological samples for electron microscopy and chemical imaging.

[53] Johnson, M. D. (2011). The acquisition of phototrophy: adaptive strategies of hosting endosymbionts and organelles. Photosynthesis research, 107, 117–132.

[54] Yellowlees, D., Rees, T. A. V., & Leggat, W. (2008). Metabolic interactions between algal symbionts and invertebrate hosts. Plant, cell & environment, 31(5), 679–694.

[55] Decelle, J., Colin, S., & Foster, R. A. (2015). Photosymbiosis in marine planktonic protists. Marine protists: diversity and dynamics, 465-500.

[56} Stoecker, D. K., Johnson, M. D., de Vargas, C., & Not, F. (2009). Acquired phototrophy in aquatic protists. Aquatic Microbial Ecology, 57(3), 279–310.

[57] Yamada, N., Bolton, J. J., Trobajo, R., Mann, D. G., Dąbek, P., Witkowski, A., … & Kroth, P. G. (2019). Discovery of a kleptoplastic ‘dinotom’dinoflagellate and the unique nuclear dynamics of converting kleptoplastids to permanent plastids. Scientific reports, 9(1), 10474.

[58] Zhuang, X., & Jiang, L. (2019). Chloroplast degradation: multiple routes into the vacuole. Frontiers in plant science, 10, 359.

[59] Krupinska, K. (2007). Fate and activities of plastids during leaf senescence. In The structure and function of plastids (pp. 433–449). Dordrecht: Springer Netherlands.

[60] Pochic, V., Gernez, P., Zoffoli, M. L., Séchet, V., Carpentier, L., & Lacour, T. (2024). Photoacclimation in the kleptoplastidic ciliate Mesodinium rubrum and its cryptophyte prey Teleaulax amphioxeia: phenotypic variability and implications for red tide remote sensing. Journal of Plankton Research, 46(2), 100–116.\

[61] Hansen, P. J., Ojamäe, K., Berge, T., Trampe, E. C., Nielsen, L. T., Lips, I., & Kühl, M. (2016). Photoregulation in a kleptochloroplastidic dinoflagellate, Dinophysis acuta. Frontiers in Microbiology, 7, 785.

[62] Kirillov, A., Mintun, E., Ravi, N., Mao, H., Rolland, C., Gustafson, L., … & Girshick, R. (2023). Segment anything. In Proceedings of the IEEE/CVF International Conference on Computer Vision (pp. 4015–4026).

[63] Gambarotto, D., Zwettler, F. U., Le Guennec, M., Schmidt-Cernohorska, M., Fortun, D., Borgers, S., … & Guichard, P. (2019). Imaging cellular ultrastructures using expansion microscopy (U-ExM). Nature methods, 16(1), 71–74.

[64] Shah, H., Olivetta, M., Bhickta, C., Ronchi, P., Trupinić, M., Tromer, E. C., … & Dey, G. (2024). Life-cycle-coupled evolution of mitosis in close relatives of animals. Nature, 1–7.

[65] Felix Mikus, Armando Rubio Ramos, Hiral Shah, Marine Olivetta, Susanne Borgers, Jonas Hellgoth, Clémence Saint-Donat, Margarida Araújo, Chandni Bhickta, Paulina Cherek, Jone Bilbao, Estibalitz Txurruka, Nikolaus Leisch, Yannick Schwab, Filip Husnik, Sergio Seoane, Ian Probert, Paul Guichard, Virginie Hamel, Gautam Dey, Omaya Dudin (2024) Charting the landscape of cytoskeletal diversity in microbial eukaryotes. bioRxiv 2024.10.18.618984; doi: 10.1101/2024.10.18.618984

[66] Catacora Grundy, A., Chevalier, F., Yee, D., LeKieffre, C., Schieber, N., Schwab, Y., … & Decelle, J. (2023). Sweet and fatty symbionts: photosynthetic productivity and carbon storage boosted in microalgae within a host. bioRxiv, 2023–12.

